# A Specialized CD107a⁺ Macrophage Subset Drives Hyperphagocytosis of Mycobacteria

**DOI:** 10.1101/2025.10.31.685620

**Authors:** Carolina Eto, Gabriela Luiz, Hernandez M. Silva, Eduarda L. Munari, Daniel A. G. B. Mendes, Bianca K. Beck, Lucas Z. Mascarin, Marick Starick, Maria Cecília C. Canesso, Bernardo S. Reis, Guilherme Silveira, Juliano Bordignon, Yonne K. T. de Menezes, Daniel S. Mansur, Edroaldo Lummertz da Rocha, André Luiz Barbosa Báfica

## Abstract

Macrophages are essential for pathogen clearance, yet phagocytic specialization among subsets is poorly defined. Bone marrow-derived macrophages cultured with L929 supernatant or M-CSF separate into FSC^lo^SSC^lo^F4/80^lo^CD11b^lo^ (FSC^lo^SSC^lo^) and FSC^hi^SSC^hi^F4/80^hi^CD11b^hi^ (FSC^hi^SSC^hi^) subsets. Transcriptomic and functional analyses reveal that FSC^lo^SSC^lo^ cells possess a hyperphagocytic program driven by enhanced actin cytoskeleton regulators (e.g., Arp2/3) and pro-inflammatory signaling (NF-κB). These cells excel at internalizing *Mycobacterium tuberculosis* virulent H37Rv and BCG through actin-mediated, cytochalasin D-sensitive mechanisms. High surface CD107a (LAMP1) expression marks this hyperphagocytic subset and correlates strongly with mycobacterial uptake. FSC^lo^SSC^lo^ macrophages produce more TNF and IL-6 upon mycobacterial or TLR2/TLR5 stimulation yet retain IFN-γ-mediated killing capacity. In vivo, CD107a⁺ alveolar macrophages in BCG-infected lungs preferentially capture bacilli and upregulate CD195, recapitulating the in vitro phenotype. These findings establish CD107a as a key surface marker of a hyperphagocytic macrophage subset and highlight its role in selective mycobacterial phagocytosis, providing new mechanistic understanding relevant to tuberculosis host defense and therapeutic development.

**Highlights:**

- Bone marrow-derived FSC^lo^SSC^lo^F4/80^lo^CD11b^lo^ macrophage subset excels at *M. tuberculosis* and BCG uptake
- Hyperphagocytic macrophages mount rapid pro-inflammatory TNF/IL-6 responses
- CD107a (LAMP1) identifies a pre-existing hyperphagocytic macrophage subset in vitro and in vivo
- BCG infection triggers CD195 (CCR5) upregulation selectively on CD107a⁺ alveolar macrophages

## Introduction

Macrophages, critical immune cells orchestrating pathogen clearance, inflammation, and tissue homeostasis, exhibit remarkable diversity in ontogeny, phenotype, and function, a concept rooted in Metchnikoff’s foundational work on phagocytosis (Tauber, 2003; Gordon and Pludemann, 2017; Ginhoux and Guilliams, 2018). Since the 1970s, bone marrow-derived macrophages (BMMs) cultured with L929-conditioned medium, a potent source of macrophage colony-stimulating factor (M-CSF), have been a cornerstone of macrophage research, enabling the generation of large, reproducible populations (Austin et al., 1971; Stanley et al., 1978; Heap et al., 2021). Flow cytometry-based experiments have identified two distinct L929-generated BMM subsets: FSC^hi^SSC^hi^ macrophages, characterized by high F4/80 and CD11b expression, and FSC^lo^SSC^lo^ macrophages, marked by lower levels of these markers (Zajd et al., 2020). However, the functional specialization of these populations, particularly their phagocytic and pathogen-specific roles, remains largely unexplored, presenting an opportunity to advance our understanding of macrophage heterogeneity.

Phagocytosis, a keystone of macrophage-mediated immunity, is a complex, multi-step process, with pathogens like *Mycobacterium tuberculosis* exploiting these cells for survival and replication. During phagocytosis, macrophages recognize *M. tuberculosis* H37Rv through pattern recognition receptors, such as Dectin-1, Toll-like receptor 2 (TLR2) and mannose receptor (CD206), which bind bacterial cell wall components like lipoarabinomannan (Ernst, 1998; Schlesinger, 1993; Rothfuchs et al, 2007). This recognition triggers actin polymerization, driven by Arp2/3 complex and Rho GTPases, to engulf the bacteria into a phagosome, a process sensitive to inhibitors like cytochalasin D (Mullins et al, 1998; Aderem and Underhill, 1999, Hinman et al, 2021; Schlam et al, 2015). Subsequently, phagosome maturation involves fusion with lysosomes, mediated by Rab GTPases, to form a phagolysosome where antimicrobial effectors attempt bacterial killing, though *M. tuberculosis* often evades destruction by inhibiting phagosome-lysosome fusion (Flannagan et al., 2012; Pieters, 2008).

In routine phagocytosis assays using L929-generated BMMs, we serendipitously observed that FSC^lo^SSC^lo^ macrophages exhibited superior uptake of *M. tuberculosis* virulent H37Rv and *M. bovis* BCG compared to FSC^hi^SSC^hi^ counterparts, challenging assumptions of functional uniformity by these cell cultures. This discovery prompted us to integrate transcriptomic profiling, phenotypic characterization, and functional assays comparing these subsets. Our results revealed that elevated surface expression of CD107a (LAMP1) and CD195 (CCR5) distinguishes hyperphagocytic macrophages with exceptional pathogen engulfment capacity. In vivo, these markers also identified a parallel subset among alveolar macrophages in BCG-infected mice. This work uncovers a novel facet of macrophage biology, offering critical insights into host-mycobacteria interactions.

## Results

### 1. Distinct actin and phagocytosis gene signatures define FSC^lo^SSC^lo^F4/80^lo^CD11b^lo^ and FSC^hi^SSC^hi^F4/80^hi^CD11b^hi^ macrophages

Building on prior studies (Zadj et al., 2020), we identified two L929-generated BMM subsets by forward scatter (FSC) and side scatter (SSC) profiles (Fig. 1A). Macrophages were categorized as FSC^lo^SSC^lo^ (∼20%, range 11–45%), with low F4/80 and CD11b expression (Fig. 1B–D), and FSC^hi^SSC^hi^ cells (∼80%, range 51–87%), with high surface F4/80 and CD11b expression (Fig. 1B–D). Similar subsets emerged in BM cell cultures treated with exogenous M-CSF, a key differentiation factor present in L929-conditioned media (Heap et al., 2021) (Fig. 1E–H). Flow cytometry-based sorting (Fig. 1I) and microscopy (Fig. 1J) revealed smaller cell areas for FSC^lo^SSC^lo^ (∼19 μm²) than FSC^hi^SSC^hi^ macrophages (∼40 μm²) (Fig. 1K), confirmed by FSC-based volume estimates (Fig. 1L). Thus, differences observed in FSC/SSC (Fig. 1A) reflect reduced forward scatter (cell size) and side scatter (internal complexity/granularity) in the FSC^lo^SSC^lo^ subset. Strikingly, bulk mRNA sequencing of sorted subsets highlighted distinct transcriptional profiles. For instance, gene ontology (GO) analysis showed FSC^hi^SSC^hi^ macrophages to be uniquely enriched in pathways such as autophagy, reactive oxygen species metabolism, response to nutrient levels, complement activation, cell recognition, lysosomal and vacuolar transport (Fig. 1M). In contrast, FSC^lo^SSC^lo^ macrophages favored actin cytoskeleton organization, response to virus, cytokine signaling, and cellular extravasation (Fig. 1M). Both subsets shared core macrophage functions such as antigen presentation, entry into host, phagocytosis, cell chemotaxis and leukocyte migration (Fig. 1M), but FSC^lo^SSC^lo^ macrophages exhibited greater enrichment in these processes. Unique sets of phagocytosis- and actin cytoskeleton-related genes distinguished the FSC^lo^SSC^lo^ and FSC^hi^SSC^hi^ subpopulations (Fig. 1N). The FSC^lo^SSC^lo^ subset was enriched for specific regulators (e.g., Arp2/3 complex, Pfn1, Coro1a) and receptors (e.g., Mincle) associated with mycobacterial uptake, consistent with its functional specialization for Mtb/BCG (Mylvaganam et al., 2021; Ferron et al, 2007; Yan et al, 2005) (Fig. 1N, Pfn1: *right panel* and Coro1a: *left panel*). These results suggest specialized actin-mediated roles in phagocytic interactions by BMM subsets.

**Figure 1.**
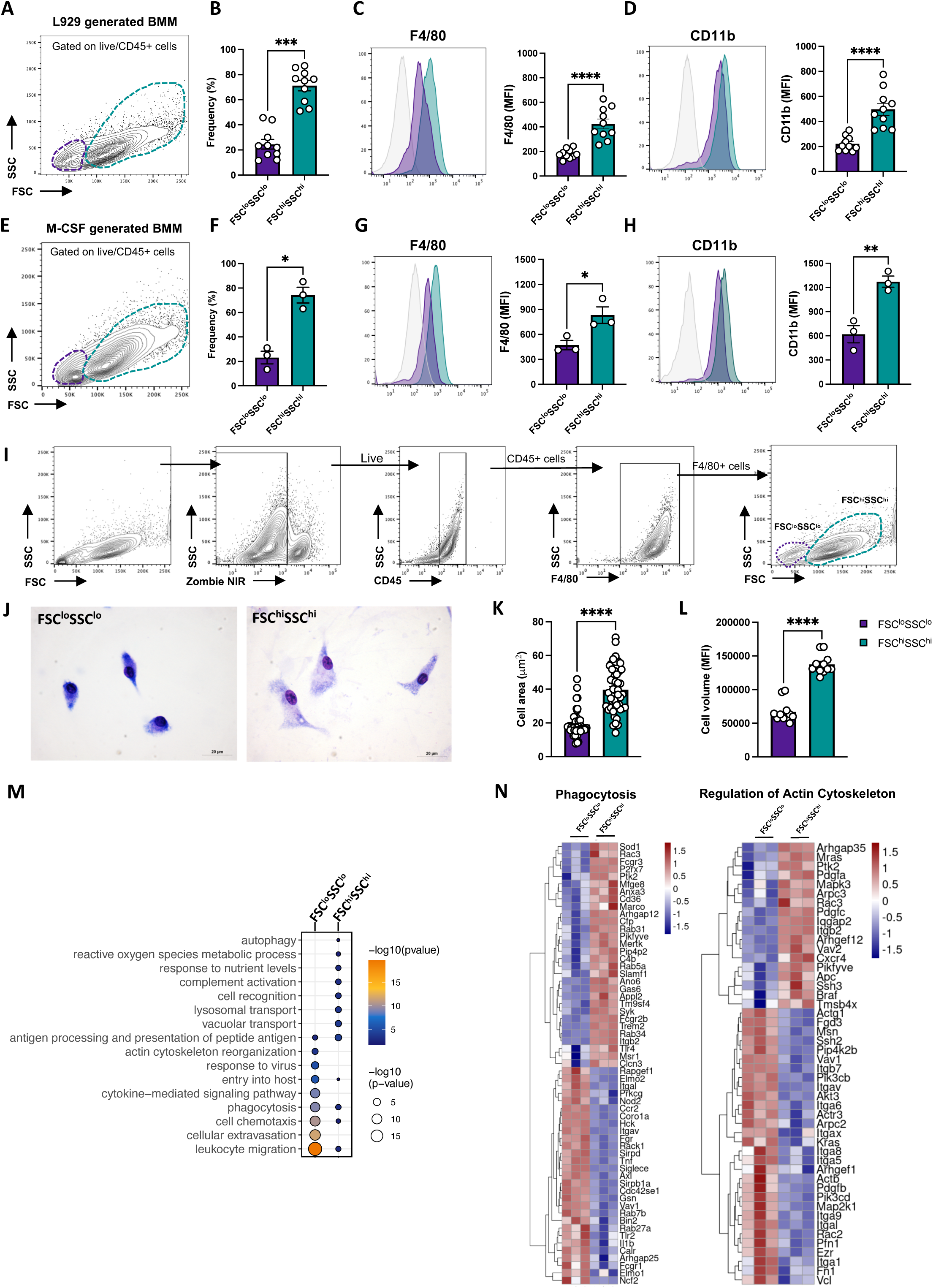
Distinct ‘Actin/Phagocytosis’ Gene Signatures in FSC^lo^SSC^lo^F4/80^lo^CD11^lo^ and FSC^hi^SSC^hi^F4/80^hi^CD11b^hi^ BM-Derived Macrophages. Bone marrow cells from C57BL/6J mice were cultured for 7 days in L929-conditioned medium or M-CSF to generate BMMs, followed by flow cytometry analysis. (A, E) Representative dot plots of forward scatter (FSC) versus side scatter (SSC) for BMMs derived using L929-conditioned medium (A) or M-CSF (E). (B, F) Frequencies of two distinct BMM populations derived with L929-conditioned medium (B) or M-CSF (F). (C, G) Representative overlaid histograms of F4/80 expression and median fluorescence intensity (MFI) on live BMMs derived using L929-conditioned medium (C) or M-CSF (G). (D, H) Representative overlaid histograms of CD11b expression and MFI on live BMMs derived with L929-conditioned medium (D) or M-CSF (H). Data in panels A-D represent mean ± SEM from ten independent experiments performed in duplicates. Data in panels E-H represent mean ± SEM from three independent experiments performed in duplicates. BM cells from C57BL/6J mice were cultured for 7 days in 20% L929-conditioned medium to generate BMMs. Subsequently, BMMs were sorted by flow cytometry, and purified F4/80^+^FSC^lo^SSC^lo^ and F4/80^+^FSC ^hi^SSC ^hi^ subpopulations were analyzed by imaging and transcriptomic profiling. (I) Gating strategy for sorting F4/80^+^FSC^lo^SSC^lo^ and F4/80^+^FSC^hi^SSC^hi^ BMMs by flow cytometry. Post-sorting analysis confirmed FSC^lo^SSC^lo^ are F4/80^lo^CD11b^lo^ and FSC^hi^SSC^hi^ are F4/80^hi^CD11b^hi^ macrophages. (J) Representative panoptic-stained images of sorted FSC^lo^SSC^lo^ and FSC^hi^SSC^hi^ BMMs at 100× magnification; scale bar = 20 μm. (K) Quantitative analysis of cell area measured using ImageJ software. (L) Quantitative analysis of cell volume based on MFI of FSC from flow cytometry. Data in K and L represent mean ± SEM of three independent experiments with 50 cells analyzed per experiment. (M) Bubble plot depicting 16 enriched Gene Ontology (GO) terms for genes upregulated in steady-state sorted FSC^lo^SSC^lo^ and FSC^hi^SSC^hi^ BMMs. (N) Heatmaps showing expression levels of differentially expressed genes in sorted FSC^lo^SSC^lo^ and FSC^hi^SSC^hi^ BMMs, with rows representing individual genes and columns representing biological replicates. Data in M and N represent mean ± SEM of three independent experiments. Statistical analyses were performed using Student’s t-test (*P < 0.05; **P < 0.01; ***P < 0.001; ****P < 0.0001).

### 2. Preferential mycobacterial phagocytosis by FSC^lo^SSC^lo^ macrophages

To investigate phagocytic interactions of BMM subsets, we performed in vitro assays using fluorescently labeled virulent H37Rv (Mtb-SYTO24) and BCG as a model system (Fig. 2A). Binding assays, adapted from established protocols (Rothfuchs et al, 2007), and quantified by flow cytometry, demonstrated that ∼75% of FSC^lo^SSC^lo^ macrophages associated with Mtb-SYTO24, compared to ∼40% of FSC^hi^SSC^hi^ macrophages, suggesting a higher binding affinity in the FSC^lo^SSC^lo^ subset (Fig. 2B). This suggests that increased initial attachment facilitates subsequent actin-mediated phagocytosis in this population, consistent with the subset’s enrichment for specific actin regulators (e.g., Arp2/3 components) and receptors that support stable adhesion and zipper-like engulfment of mycobacteria. Consistent with this concept, phagocytosis assays further revealed that FSC^lo^SSC^lo^ BMM associated with ∼70% of Mtb-SYTO24, significantly outperforming FSC^hi^SSC^hi^ macrophages, which internalized ∼30%, with elevated mean fluorescence intensity (MFI) reflecting greater bacterial loads per FSC^lo^SSC^lo^ cell (Fig. 2C). In contrast, FSC^hi^SSC^hi^ macrophages exhibited superior phagocytosis of zymosan-FITC (fungal-derived) (Fig. 2D) and *Salmonella*-SYTO24 (Gram-negative) (Fig. 2E). Corroborating flow cytometry results, fluorescence microscopy of Mtb-SYTO24-infected macrophages showed that smaller cells (area < 30 μm²) contained an average of 2.52 bacilli per cell, compared to 1.32 bacilli in larger cells (> 30 μm²) (Fig. 2F and G; p = 0.0011). Next, we compared uptake kinetics and capacity of FSC^lo^SSC^lo^ and FSC^hi^SSC^hi^ macrophages during exposure to *M. bovis* BCG. Time-course assays showed enhanced mycobacterial uptake by FSC^lo^SSC^lo^ macrophages from 1h post-infection (Fig. 2H and I). Moreover, quantitative four-parameter logistic regression of bacterial MOI (exposure) versus MFI values (uptake) confirmed greater infection burden in FSC^lo^SSC^lo^ macrophages compared to FSC^hi^SSC^hi^ cells across various MOIs (Fig. 2J; R² ∼ 0.99 for both subsets). Surprisingly, although FSC^hi^SSC^hi^ cells exhibited higher expression of classical mycobacterial phagocytosis receptors - including CD206, Marco, CD36, Mcl, and Dectin-1 (Fig. 1N, 3A) - FSC^lo^SSC^lo^ macrophages demonstrated greater bacterial uptake. These findings suggest that the FSC^lo^SSC^lo^ subset exhibits enhanced mycobacterial uptake capacity via non-canonical mechanism(s).

**Figure. 2.**
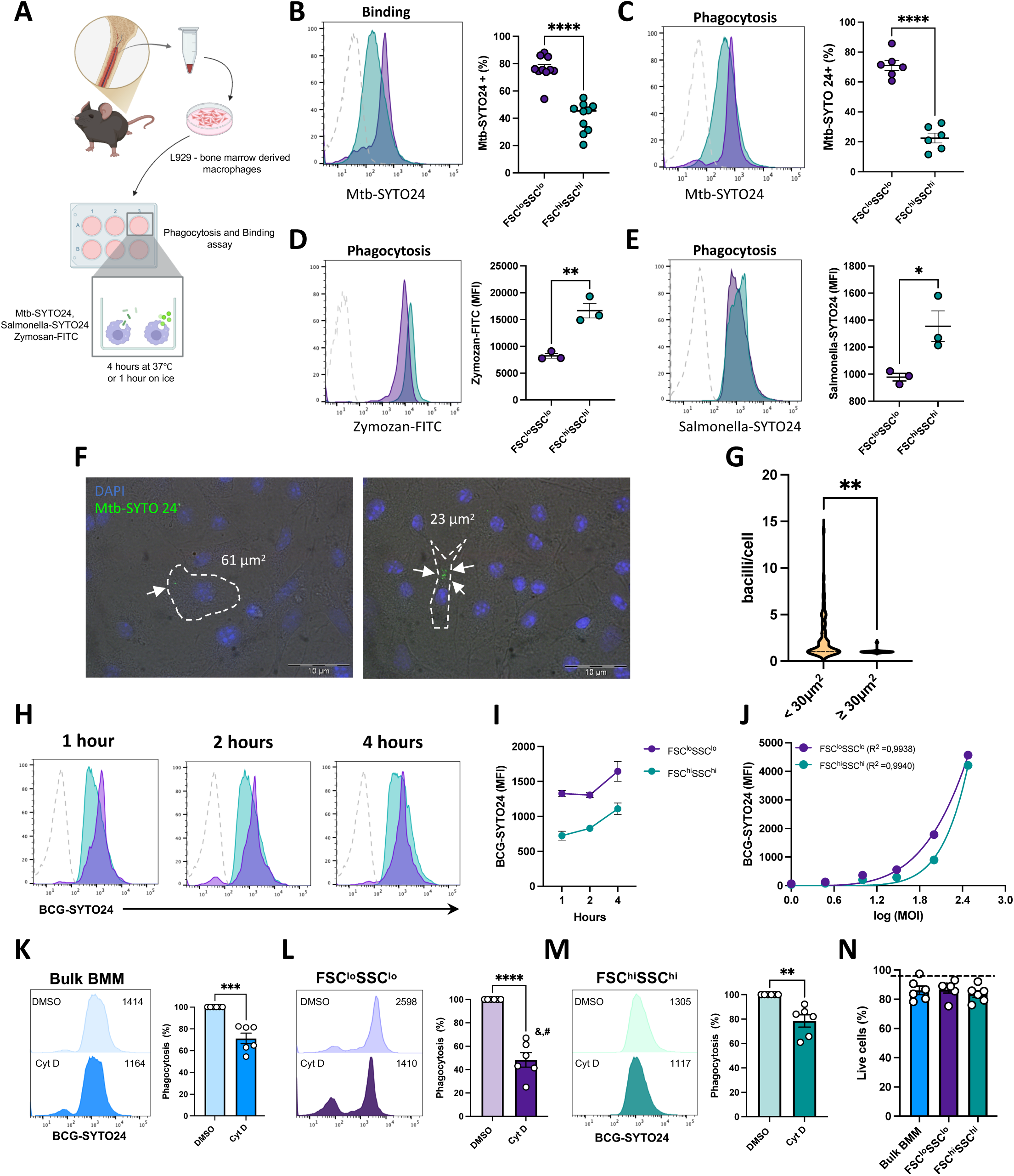
High Phagocytic Activity of BM-Derived FSC^lo^SSC^lo^ Macrophages Toward Mycobacteria. BMMs from C57BL/6J mice were exposed *M. tuberculosis* H37Rv (Mtb)-SYTO24 (MOI 10), zymosan-FITC (50 μg/mL) for 4 h at 37°C or Salmonella -SYTO24 (MOI 10) for 30 minutes at 37°C, and particle association was assessed by flow cytometry. (A) Experimental design. (B) Representative histogram and frequency of binding Mtb-SYTO24^+^ FSC^lo^SSC^lo^ (purple) and FSC^hi^SSC^hi^ (teal) macrophages. (C) Representative histogram and frequency of phagocytosed Mtb-SYTO24^+^ FSC^lo^SSC^lo^ (purple) and FSC^hi^SSC^hi^ (teal) macrophages. (D) Representative histogram and Median fluorescence intensity (MFI) of zymosan-FITC phagocytosis in SSC^lo^FSC^lo^and SSC^hi^FSC^hi^ macrophages. (E) Representative histogram and MFI of Salmonella-SYTO24 phagocytosis in SSC^lo^FSC^lo^and SSC^hi^FSC^hi^ macrophages. Data in panels B represent mean ± SEM from ten independent experiments performed in duplicates. Data in panels C represent mean ± SEM from six independent experiments performed in duplicates. Data in panels D-E represent mean ± SEM from three independent experiments performed in duplicates. BMMs exposed to Mtb-SYTO24 (MOI 10) for 4 h at 37°C were analyzed by fluorescence microscopy. (F) Representative images of two macrophages showing cell areas and fluorescent mycobacteria; cell nuclei labeled with DAPI; Mtb labeled with SYTO24; scale bar = 10 μm. (G) Violin plot of bacilli counts in 96 Mtb-SYTO24-infected macrophages, quantified from 50 fields across two independent experiments, as described in (F). BMMs were exposed to *M. bovis* BCG-SYTO24 (MOI 10) for varying time points, and phagocytosis was assessed by flow cytometry. (H) Representative histograms of BCG-SYTO24 uptake. (I) MFI of BCG-SYTO24 phagocytosis over time in FSC^lo^SSC^lo^ and FSC^hi^SSC^hi^ macrophages. Data in panels H-I represent mean ± SEM from two-four independent experiments. Flow cytometry analysis of BCG-SYTO24-exposed macrophages and gated on FSC^lo^SSC^lo^ and FSC^hi^SSC^hi^ events. (J) Data are presented as a sigmoid curve fit of six independent experiments, depicting BCG-SYTO24 multiplicity of infection (MOI, exposure) versus mean fluorescence intensity (MFI, uptake), as detailed in M&M. BMMs were pretreated with cytochalasin D (5 μg/mL) for 30 min, followed by BCG-SYTO24 exposure (MOI 10) for 4 h at 37°C, and analyzed by flow cytometry. (K) MFI of BCG-SYTO24 phagocytosis in bulk BMMs. (L) MFI of BCG-SYTO24 phagocytosis in FSC^lo^SSC^lo^ macrophages. (M) MFI of BCG-SYTO24 phagocytosis in FSC^hi^SSC^hi^ macrophages. (N) Frequency (%) of live bulk BMMs, FSC^lo^SSC^lo^, and FSC^hi^SSC^hi^ macrophages following cytochalasin D treatment. Data in panels K-N represent mean ± SEM from three independent experiments. Data were analyzed using Student’s t-test (*P < 0.05; **P < 0.01; ***P < 0.001; ****P < 0.0001) or one-way ANOVA (#P < 0.05; &P < 0.01 for Cytochalasin D-treated bulk BMMs vs. FSC^lo^SSC^lo^ and FSC^hi^SSC^hi^ vs. FSC^lo^SSC^lo^ macrophages, respectively).

Next, we investigated the phagocytic mechanisms of FSC^lo^SSC^lo^ and FSC^hi^SSC^hi^ BMMs, which showed distinct gene expression profiles for phagocytosis and actin cytoskeleton regulation (Fig. 1N). We tested whether actin-mediated processes underpin their differential uptake of mycobacteria by treating cells with cytochalasin D, an actin polymerization inhibitor, and quantifying phagocytosis via flow cytometry, as established previously (Hinman et al., 2021). In bulk BMMs, cytochalasin D reduced mycobacterial uptake by 28% (range: 18–46%, p < 0.001, treated vs. vehicle; Fig. 2K). Notably, FSC^lo^SSC^lo^ macrophages exhibited greater sensitivity, with a 49% reduction in phagocytosis (range: 33–75%, p < 0.0001, treated vs. vehicle; Fig. 2L) compared to a 21.7% reduction in FSC^hi^SSC^hi^ macrophages (range: 8–39%, p < 0.01, treated vs. vehicle; Fig. 2M). Although FSC^lo^SSC^lo^ macrophages exhibited superior phagocytosis of mycobacteria (Fig. 2B, C, H, I and Supplementary Fig. 1A), this phenotype was accompanied by a broader increase in actin dependence. Parallel dose–response experiments demonstrated that the FSC^lo^SSC^lo^ subset also displayed greater sensitivity to cytochalasin D during internalization of *Salmonella* and zymosan particles compared with the FSC^hi^SSC^hi^ subset (Supplementary Fig. 1A, B), without loss of cell viability (Fig. 2N). These results suggest that the increased reliance on cytochalasin D-sensitive, actin-mediated mechanisms in FSC^lo^SSC^lo^ macrophages reflects a general feature of their phagocytic program rather than a pathogen-specific trait.

### 3. Differential TLR sensing and cytokine responses by FSC^lo^SSC^lo^F4/80^lo^CD11b^lo^ macrophages

Transcriptomics revealed enriched innate sensing genes (e.g., *Tlr1*, *Tlr2* and *Tlr5*) in FSC^lo^SSC^lo^ macrophages, while FSC^hi^SSC^hi^ cells favored *Clec7a* (Dectin-1) and *Tlr4* (Fig. 3A). These transcriptomics findings were further validated by exposing bulk BMMs to known TLR agonists in vitro and measuring intracellular TNF accumulation via flow cytometry (Fig. 3B). TLR agonist stimulation confirmed FSC^lo^SSC^lo^ macrophages as primary TNF producers for Pam3Cys, HK-BCG, and Flagellin, whereas LPS triggered TNF in FSC^hi^SSC^hi^ cells (Fig. 3B). Enhanced *Clec7a* expression aligned with increased phagocytosis of zymosan, a Dectin-1 ligand (Brown et al., 2003), by FSC^hi^SSC^hi^ macrophages (Fig. 2D). FSC^lo^SSC^lo^ macrophages showed pro-inflammatory gene enrichment (TNF, IL-6), while FSC^hi^SSC^hi^ cells leaned toward IL-10 signaling (Fig. 3C). Next, we used mycobacteria to investigate the kinetics of cytokine secretion by flow cytometry-based cell sorting purified FSC^hi^SSC^hi^ and FSC^lo^SSC^lo^ macrophages. Culture supernatants from sorted FSC^lo^SSC^lo^ macrophages at 24h and 48h post-Mtb exposure showed significantly higher TNF levels and lower IL-10 levels compared to FSC^hi^SSC^hi^ macrophages (Fig. 3D). For IL-6 production, FSC^lo^SSC^lo^ macrophages exhibited elevated levels at 24h post-infection, whereas FSC^hi^SSC^hi^ macrophages peaked at 48h (Fig. 3D). Despite higher Mtb uptake by FSC^lo^SSC^lo^ cells (Fig. 3E), both subsets exhibited similar intracellular pathogen growth (Fig. 3F) and IFN-γ-induced killing at 24h and 48h (Fig. 3G and I), with comparable nitrite production (Fig. 3H and J). Collectively, these findings indicate that FSC^lo^SSC^lo^ macrophages strongly respond to TLR2 and TLR5 ligands and maintain IFN-γ-mediated mycobacterial killing.

**Figure 3.**
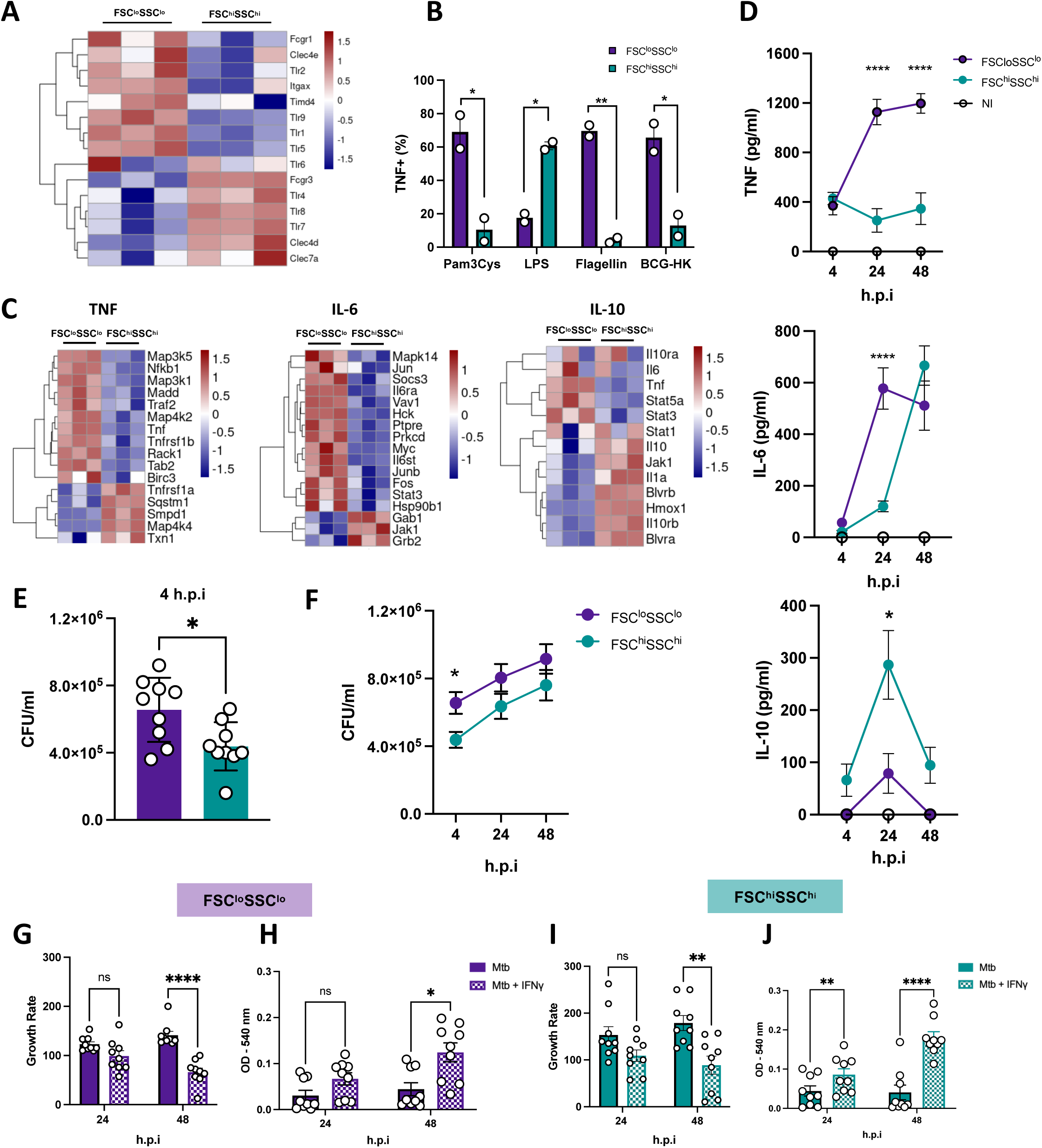
Differential TLR Recognition and Cytokine Production in BM-Derived FSC^lo^SSC^lo^ Macrophages. (A) Heatmap of differentially expressed innate immune receptor genes in sorted FSC^lo^SSC^lo^ and FSC^hi^SSC^hi^ BMMs from C57BL/6J mice, as described in Fig. 1I. Bulk BMMs were stimulated with Pam3CSK4 (100 ng/mL), *Escherichia coli* LPS (100 ng/mL), flagellin (100 ng/mL), for 2 h or heat-killed BCG (MOI 10) for 4 h at 37°C, and intracellular TNF was quantified by flow cytometry. (B) Percentage of TNF-producing FSC^lo^SSC^lo^ and FSC^hi^SSC^hi^ macrophages following TLR ligand stimulation. (C) Heatmaps depicting expression levels of differentially expressed genes in TNF, IL-6, and IL-10 signaling pathways in sorted FSC^lo^SSC^lo^ and FSC^hi^SSC^hi^. Sorted FSC^lo^SSC^lo^F4/80^lo^CD11b^lo^ and FSC^hi^SSC^hi^F4/80^hi^CD11b^hi^ BMMs (as in Fig. 1I) were exposed to Mtb (MOI 3) at 4 h, 24 h and 48 h. Culture supernatants were analyzed for TNF, IL-6, and IL-10 by ELISA. (D) Cytokine production by sorted FSC^lo^SSC^lo^ and FSC^hi^SSC^hi^ macrophages post-Mtb infection. Sorted FSC^lo^SSC^lo^F4/80^lo^CD11b^lo^ and FSC^hi^SSC^hi^F4/80^hi^CD11b^hi^ BMMs were exposed to Mtb (MOI 3) at different time points, and (E) phagocytosis and (F) intracellular bacterial growth were assessed by colony-forming unit (CFU) counts. (G, I) Mycobacterial growth rates at 24 and 48 h, calculated as ratios relative to the 4 h time point, in FSC^lo^SSC^lo^ and FSC^hi^SSC^hi^ macrophages, with or without IFN-γ (100 UI/mL). (H, J) Nitrite levels in culture supernatants of FSC^lo^SSC^lo^ and FSC^hi^SSC^hi^ macrophages from panels G and I. Data represent the mean ± SEM of two (B) or three (A, C, D–J) independent experiments. Statistical analyses were conducted using Student’s t-test (E and B) or two-way ANOVA with Tukey’s multiple comparisons test (D, F-J). *P < 0.05; **P < 0.01; ***P < 0.001; ****P < 0.0001.

### 4. Surface CD107a and CD195 in FSC^lo^SSC^lo^ macrophages mark enhanced mycobacterial uptake in vitro

To further characterize macrophage subset phenotypes, we conducted flow cytometry using a comprehensive panel of surface markers, aiming to identify markers associated with enhanced mycobacterial uptake in L929-induced BMMs. FSC^hi^SSC^hi^ macrophages expressed higher levels of CD45, CD64, CD74, CD88, CD115, CD206, and CD207, while FSC^lo^SSC^lo^ macrophages showed elevated CD107a (LAMP1) and CD195 (CCR5) (Fig. 4A–B), consistent across M-CSF- and L929-derived BMMs. Next, we assessed marker dynamics during BCG infection. While CD107a expression remained unchanged in both subsets, CD195 increased ∼5-fold in FSC^lo^SSC^lo^ macrophages versus ∼2-fold in FSC^hi^SSC^hi^ cells (Fig. 4C–D) following BCG exposure. Notably, in infected FSC^lo^SSC^lo^ macrophages, the CD107a^+^ subset exhibited even higher mycobacterial loads compared to their CD107a^−^ counterparts (Fig. 4E). Surface CD107a levels remained high and stable post-phagocytosis in time-course analyses (data not shown), suggesting its role as a marker rather than an uptake mediator. Furthermore, flow cytometry analysis of infected macrophages from 6 independent experiments revealed a strong positive monotonic correlation between BCG and CD107a MFI values (Spearman’s ρ ∼ 1.0, p = 0.0028), and a moderate correlation between BCG and CD195 MFI values (Spearman’s ρ ∼ 0.83, p = 0.058) in FSC^lo^SSC^lo^ macrophages (Fig. 4F). Given its reported role in binding mycobacterial molecules in human dendritic cells (Floto et al., 2006), we investigated the potential involvement of CD195 in mycobacterial uptake by FSC^lo^SSC^lo^ macrophages using BMMs from CD195-deficient mice. Flow cytometry after 4h Mtb exposure showed CD195-deficient FSC^lo^SSC^lo^ macrophages retained bacterial phagocytosis (Fig. 4G–I). Although CD206 expression was elevated in FSC^hi^SSC^hi^ macrophages (Fig. 4A), its lower expression in FSC^lo^SSC^lo^ cells suggests that this mannose receptor is unlikely to contribute to their selective mycobacterial binding/uptake, pointing to alternative receptors or mechanisms. Nevertheless, our in vitro data suggest CD107a and CD195 may serve as surface markers for identifying macrophages with heightened phagocytic activity toward mycobacteria.

**Figure 4.**
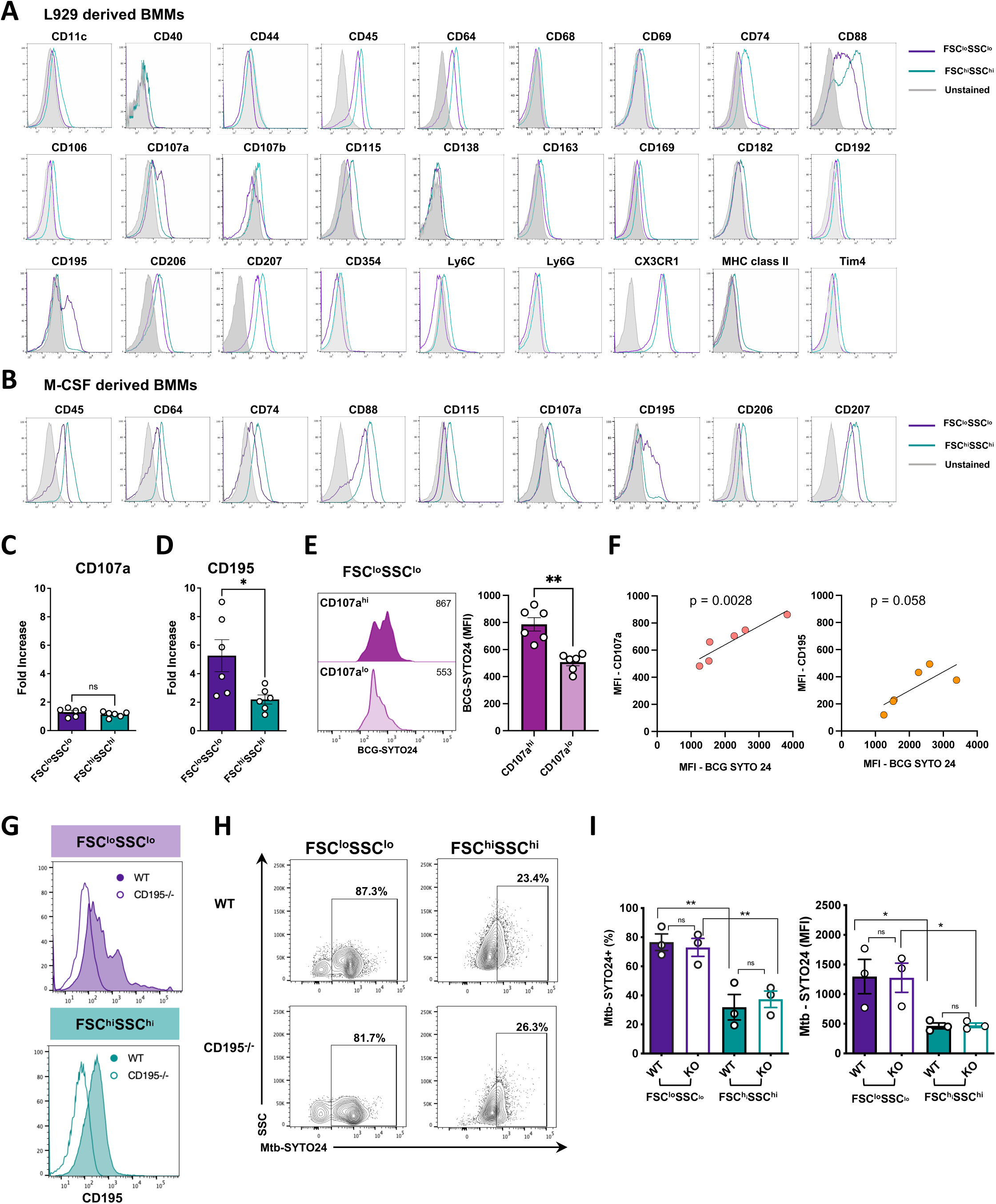
Surface expression of CD107a and CD195 in FSC^lo^SSC^lo^ and FSC^hi^SSC^hi^ macrophage subsets. Bone marrow cells from C57BL/6J mice were cultured for 7 days in (A) L929-conditioned medium or (B) M-CSF to generate BMMs, followed by flow cytometry analysis. (A) Representative histograms of surface molecules expressed in FSC^lo^SSC^lo^ (purple) and FSC^hi^SSC^hi^ (teal) BMMs derived with L929-conditioned medium or M-CSF (B). BMMs were exposed to BCG (MOI 10) for 4 h, and CD107a and CD195 expression was assessed by flow cytometry. Fold increase of median fluorescence intensity (MFI) of surface (C) CD107a and (D) CD195 over uninfected cells in FSC^lo^SSC^lo^ and FSC^hi^SSC^hi^ BMMs. FSC^lo^SSC^lo^ cells from experiments described in (C and D), were gated on BCG^+^ events and analyzed for expression of CD107a. (E) Comparison of BCG-SYTO24 MFI between CD107a^hi^ (dark purple) and CD107a^lo/-^(light purple) infected FSC^lo^SSC^lo^ macrophages. (F) Scatter plots showing correlations between BCG MFI and surface markers in FSC^lo^SSC^lo^ macrophages at 4 h post-BCG exposure from 6 independent experiments. Left: Correlation between BCG and CD107a MFI (Spearman’s ρ = 1.0, p = 0.0028, n = 6). Right: Correlation between BCG and CD195 MFI (Spearman’s ρ ∼ 0.83, p = 0.058, n = 6). Points represent MFI values, plotted on linear axes. BMMs from C57BL/6J and CD195^−/−^ mice were exposed to Mtb-SYTO24 (MOI 10) for 4 h, and analyzed by flow cytometry. (G) Representative histograms confirming CD195 deficiency in CD195^−/−^ BMMs. (H) Dot plots showing frequency of Mtb-SYTO24-infected WT and CD195^−/−^ FSC^lo^SSC^lo^ and FSC^hi^SSC^hi^ macrophages. (I) Frequency and MFI of Mtb-SYTO24-exposed WT and CD195^−/−^ FSC^lo^SSC^lo^ and FSC^hi^SSC^hi^ macrophages. Data represent mean ± SEM of three (panels C-E and G-I) and six-nine (panel F) independent experiments. Statistical analyses were performed using Student’s t-test (C-E and I). *P < 0.05; **P < 0.01.

### 5. CD107a identify hyperphagocytic AMs in vivo

To investigate whether these surface molecules serve as markers for hyperphagocytic activity by macrophages in vivo, we examined lung macrophages in C57BL/6J mice using an aerogenic BCG infection model (Cohen et al., 2018). Lung macrophages were categorized as alveolar macrophages (AM, LiveCD45^+^CD64^+^F4/80^+^CD11b^lo^SiglecF^hi^CD11c^hi^) or interstitial macrophages (IM, LiveCD45^+^CD64^+^F4/80^+^CD11b^hi^SiglecF^−^CD11c^lo^) based on established flow cytometry panels (Fig. 5A) (Bedoret et al., 2009; Misharin et al., 2013; Pisu et al., 2021; Menezes et al., 2024). Consistent with prior findings (Cohen et al., 2018), AMs displayed significantly greater association with fluorescent BCG bacilli compared to IMs at 18h post-infection, as assessed by flow cytometry (Fig. 5B and C). We then quantified surface expression of CD107a and CD195 on AMs and IMs from saline- and BCG-exposed mice. AMs exhibited higher baseline expression of both markers compared to IMs (Fig. 5D and E). Post-infection, CD107a levels remained unaltered in AMs, aligning with in vitro observations (Fig. 4C), whereas CD195 expression increased approximately 6-fold in AMs but not IMs (Fig. 5F). As CD107a expression remains unchanged during BCG infection, it may serve as a reliable surface marker for AMs with enhanced mycobacterial uptake capacity. In support of this hypothesis, we gated on AMs (LiveCD45^+^CD64^+^F4/80^+^CD11b^lo^SiglecF^hi^CD11c^hi^) and, based on CD107a expression, we identified two AM subsets: CD107a^+^ and CD107a^−^, each comprising ∼50% of AMs (Fig. 5G). CD107a^+^ and CD107a^−^ displayed similar FSC and SSC profiles (data not shown). Notably, the CD107a^+^ subset showed significantly higher frequencies of BCG-SYTO24^+^ macrophages and elevated MFI for BCG uptake (Fig. 5H-J). Our results demonstrate that CD107a identifies a hyperphagocytic subset of AMs in vivo with enhanced BCG uptake. Following BCG exposure, CD107a^+^ AMs significantly upregulate surface CD195 expression, highlighting their role in mycobacterial interactions and suggesting CD107a and CD195 as key markers of hyperphagocytic macrophages in mycobacteria infection.

**Figure 5.**
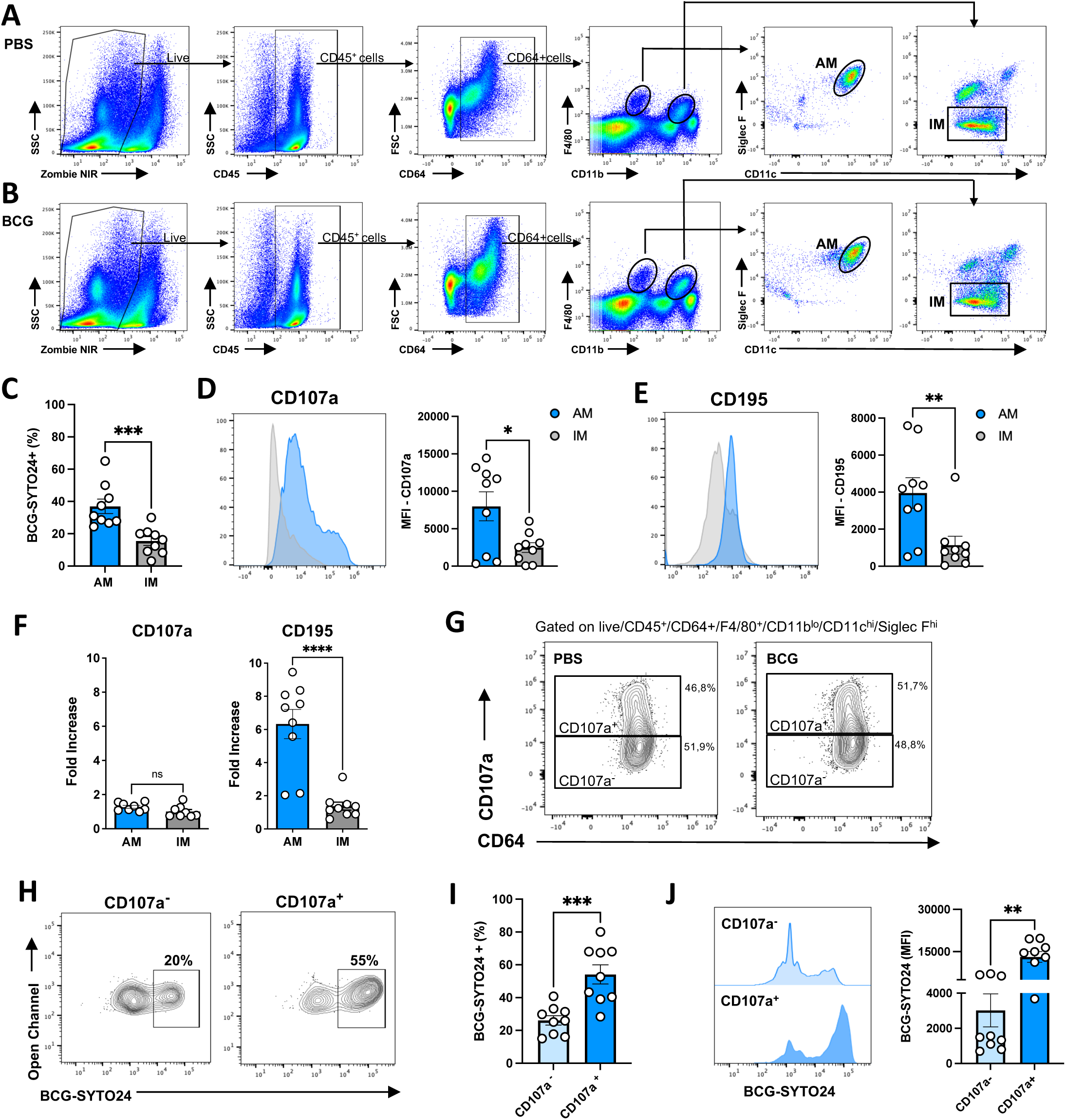
CD107a and CD195 expression in alveolar macrophages is associated with enhanced mycobacterial phagocytosis in vivo. Lung cell preparations from C57BL/6J mice exposed intratracheally (i.t.) to saline or *M. bovis* BCG (10^8^ CFU/mouse) were analyzed by flow cytometry (A and B), as described in the Materials and Methods. (A) Gating strategy for identifying alveolar (AM; CD45^+^CD64^+^F4/80^+^CD11b^lo^SiglecF^hi^CD11c^hi^) and interstitial (IM; CD45^+^CD64^+^F4/80^+^CD11b^hi^SiglecF^neg^CD11c^lo^) macrophage populations in flow cytometry data from total lung cell preparations of saline-exposed mice. (C) Frequency of BCG-SYTO24^+^AM and IM. Representative histogram and MFI of CD107a (D) and CD195 (E) expression in AM and IM from saline-exposed mice. (F) Fold increase in surface receptor expression, showing significant CD195 upregulation in AM but not IM from BCG-exposed mice compared to saline-exposed controls. (G) Representative contour dot plot of AM showing CD107a versus CD64 expression frequencies in C57BL/6J mice exposed to saline or BCG for 18 h, analyzed by flow cytometry. (H) Representative contour dot plot of BCG^+^ cells in CD107a^+^ or CD107a^−^ AMs. (I) BCG-SYTO24^+^ frequencies and (J) MFI from CD107a^−^ and CD107a^+^ gates (as described in H), analyzed at 18 h post-BCG infection. Data pooled from four independent experiments (n = 9 mice/group). Statistical analyses were performed using Student’s t test. P-value **<0.01; ***<0.001; ****<0.0001.

## Discussion

Macrophage heterogeneity prompts questions about subset-specific roles in pathogen interactions. We identified a hyperphagocytic FSC^lo^SSC^lo^(F4/80^lo^CD11b^lo^) BMM subset with elevated CD107a and CD195 expression, indicating enhanced mycobacterial uptake (Fig. 2C–E). In vivo, CD107a^+^ AMs in BCG-infected mice showed higher mycobacterial loads, mirroring in vitro findings. These results suggest CD107a and, upregulation of CD195, mark hyperphagocytic macrophages, with potential implications for TB host defense strategies.

Transcriptomic analysis of FSC^lo^SSC^lo^ BMMs reveals enriched phagocytosis, actin cytoskeleton, including coronin 1a and fibronectin 1 as well as pro-inflammatory cytokine pathways (Fig. 1N; Pasula et al., 2002; Jayachandran et al., 2007). Our transcriptomic analysis revealing enriched phagocytosis, actin cytoskeleton, and innate sensing pathways in macrophage subsets was substantiated by functional assays, including TNF and IL-6 production in response to TLR2, TLR4 and TLR5 ligands (Fig. 3B) and elevated cytokine secretion post-Mtb infection (Fig. 3D), confirming the pro-inflammatory and phagocytic specialization of each subset. Additional studies are required to further characterize the specific roles of enriched genes linked to pathogen recognition and internalization, such as Tlr2 and Pfn1, in driving the selective mycobacterial uptake by FSC^lo^SSC^lo^ macrophages. While both macrophage subsets exhibited high expression of phagocytic receptors, they demonstrated distinct transcriptional programs associated with phagocytosis, suggesting previously unknown entry mechanisms utilized by mycobacteria.

While virulent *M. tuberculosis* strains (e.g., H37Rv) exhibit enhanced immune evasion compared to BCG (e.g., via ESX-1-mediated phagosomal perforation), core mechanisms of initial macrophage uptake - including actin-mediated phagocytosis and primary engagement of complement receptors - are conserved across attenuated (BCG, H37Ra) and virulent strains (H37Rv, Erdman). For example, studies have shown that both virulent and attenuated strains utilize complement receptors as the dominant pathway for entry, with virulent strains additionally recruiting mannose receptors in some contexts (Schlesinger et al., 1993, J Immunol). In vivo, BCG infection reliably models alveolar macrophage uptake and recruitment of phagocytic subsets, mirroring key early events in TB pathogenesis.

The superior mycobacterial uptake by the FSC^lo^SSC^lo^ subset is closely associated with their enhanced binding capacity, indicating that stronger initial surface association may actively promote phagocytosis in this population. This is consistent with known mechanisms of mycobacterial entry, where stable receptor-mediated adhesion (e.g., via complement and mannose receptors) is a prerequisite for efficient actin polymerization and phagocytic cup closure. While flow-based assays cannot fully decouple binding from internalization without orthogonal controls (e.g., confocal Z-stack analysis), our data support a model in which enhanced adhesion in the FSC^lo^SSC^lo^ subset drives their mycobacteria-selective hyperphagocytosis.

Although the FSC^lo^SSC^lo^ subset represents only ∼20% of BMMs, it accounts for the majority of mycobacterial internalization. This phenotype is partially reduced by cytochalasin D, suggesting that actin polymerization is central to the hyperphagocytic behavior of FSC^lo^SSC^lo^ BMMs (Fig. 2J–M). This partial sensitivity aligns with the complex, multi-receptor nature of mycobacterial phagocytosis, which can involve partial actin-independent components, differing from stronger inhibition reported for Gram-positive cocci (Kapetanovic et al., 2007; Ribes et al., 2010) or zymosan (this study). Their heightened sensitivity to actin disruption (Fig. 2K–L) may relate to the enriched expression of actin cytoskeleton regulators such as *Pfn1* and *Coro1a* (Fig. 1N), suggesting distinct actin dynamics in this subset. Consistent with a broader role for actin polymerization, cytochalasin D also reduced phagocytosis of fluorescently labeled zymosan or *Salmonella* particles. However, FSC^lo^SSC^lo^ macrophages displayed comparatively lower uptake of these particles than FSC^hi^SSC^hi^ cells (Fig. 2B–C), suggesting that their dominance in mycobacterial uptake is unlikely to reflect a generalized increase in phagocytic capacity. We speculate that upstream recognition or targeting mechanisms may contribute to the selective engagement of mycobacteria by this subset.

Transcriptomic enrichment for Gene Ontology terms related to “phagocytosis” and “actin cytoskeleton reorganization” in the FSC^lo^SSC^lo^ subset (Fig. 1M–N) therefore reflects a distinct functional program rather than indiscriminate hyperphagocytosis. Functional assays confirmed preferential uptake of mycobacteria (Mtb/BCG) by FSC^lo^SSC^lo^ cells, whereas *Salmonella* and zymosan were more efficiently internalized by FSC^hi^SSC^hi^ macrophages, which express higher levels of complementary receptors such as *Marco*, *Msr1*, and *Cd36* (Fig. 1N). These patterns suggest subset-specific phagocytic specialization: FSC^lo^SSC^lo^ macrophages appear optimized for actin-mediated handling of mycobacteria, potentially involving Arp2/3-associated mechanisms and C-type lectins, while FSC^hi^SSC^hi^ cells preferentially engage pathways suited for other microbial particles. Notably, the transcriptomic enrichment of actin regulators in FSC^lo^SSC^lo^ macrophages is correlative, and direct functional studies will be required to establish their specific roles in driving this phenotype. Elevated *Tlr2* expression in FSC^lo^SSC^lo^ macrophages fuels robust TNF and IL-6 with low IL-10 production, suggesting a pro-inflammatory environment that mycobacteria might exploit (Underhill et al., 1999, Drennan et al., 2004, Bafica et al, 2005). The earlier peak of TNF and IL-6 production by FSC^lo^SSC^lo^ macrophages at 24h post-Mtb infection, compared to the delayed IL-6 peak at 48h in FSC^hi^SSC^hi^ macrophages (Fig. 3D), suggests a rapid pro-inflammatory response that may enhance early mycobacterial control in TB, though sustained inflammation could contribute to tissue pathology if unchecked. Yet, their IFN-γ-induced mycobactericidal activity matches FSC^hi^SSC^hi^ cells (Fig. 3E), indicating intact IFN-γ-mediated effector functions. In vivo, CD107a^+^ AMs’ rapid BCG binding as early as 2h post-infection (data not shown) suggests a pre-primed hyperphagocytic state, potentially amplifying early mycobacterial interactions in the lung.

While high surface CD107a expression strongly correlates with and marks the hyperphagocytic FSC^lo^SSC^lo^ subset (Spearman’s ρ = 1.0, p = 0.0028; Fig. 4E–F), our data do not demonstrate a direct causal role for CD107a in mediating uptake. Instead, it likely reflects heightened phagosomal activity, lysosomal biogenesis, or membrane dynamics associated with superior actin-driven phagocytosis in this population. Future conditional knockouts or CRISPR screens will clarify this receptor’s role. Moreover, CD195-deficient BMMs showed no defect in mycobacterial uptake (Fig. 4H–I). This result indicates that CD195 primarily serves as a phenotypic marker of this specialized population rather than a direct functional driver or receptor mediating uptake. The specific receptor(s) responsible for the observed hyperphagocytosis remain to be identified and may involve other differentially expressed molecules (e.g., C-type lectins such as Mincle) or synergistic receptor engagement. Given the role of CD195 in cell migration, it is possible that CD107a^+^CD195^hi^ AMs may act as “Trojan horses,” facilitating mycobacterial dissemination. For instance, Kirby et al. (2009) showed that a subset of AM migrates to draining lymph nodes in a *Streptococcus pneumoniae* lung infection model. Future studies on macrophage trafficking and mechanistic roles of CD107a and CD195 are needed to further elucidate their contributions to mycobacterial interactions.

This study has several limitations. Our findings rely on mouse models, which may not fully recapitulate human macrophage biology, and short-term infection protocols that do not reflect the chronic nature of TB. Furthermore, although virulent *M. tuberculosis* H37Rv was used in key in vitro phagocytosis experiments (Fig. 2), biosafety constraints required the use of attenuated *M. bovis* BCG for in vivo validation and several functional assays. The in vivo experiments were performed at 18h post-exposure to specifically examine early phagocytic events. While this acute model highlights the enhanced phagocytic capacity of resident CD107a⁺ alveolar macrophages during initial uptake, it remains to be determined whether recruited monocyte-derived interstitial macrophages (Antonelli et al., 2010; Huang et al., 2018; Lee et al., 2020; Skold and Behar, 2008) can acquire a similar hyperphagocytic phenotype during the chronic phases of TB infection. Nevertheless, our findings define a previously unrecognized hyperphagocytic macrophage subset and establish CD107a/LAMP1 as a reliable surface marker, providing mechanistic insights into mycobacterial uptake and potential targets for intervention.

## Materials and Methods

### Ethics Statement

All animal-related protocols were approved by the Committee of Ethics on the Use of Animals from Federal University of Santa Catarina (CEUA/UFSC/#9566210722).

### Preparation of Bacteria

Frozen stocks of *Mycobacterium tuberculosis* (Mtb, H37Rv strain), *M. bovis* BCG (BCG, Danish strain), and *Salmonella enterica* (ATCC 14028) were maintained in biosafety containment facilities at Federal University of Santa Catarina. Briefly, mycobacterial cultures were grown in Middlebrook 7H9 liquid medium supplemented with OADC for 15-21 days at 37°C until reaching ideal optical density. Bacteria concentration was determined by plating onto 7H10 Middlebrook agar. *S. enterica* cultures were grown in BHI liquid medium, and bacteria concentration was determined by plating onto LB agar. For fluorescence assays, bacterial suspensions were incubated with SYTO24 (1µM) for 1 hour at 37°C, as previously described (Yamashiro et al., 2016), and then washed with sterile PBS (Gibco, 70011044) prior to infections. SYTO24 labeling, used for fluorescence-based detection of *M. tuberculosis, M. bovis* BCG, and *S. enterica*, does not alter bacterial infectivity or growth, as previously validated (Yamashiro et al., 2016; Delbogo et al., 2019). Heat-killed BCG was obtained by boiling bacterial suspension at 95°C for 10 minutes.

### Preparation of L929 cell conditioned medium

L929 murine cells were cultured in DMEM (Gibco, 11965092) supplemented with 10% heat-inactivated Fetal Bovine Serum (FBS, Gibco, 12657029), 2 mM L-glutamine (Gibco, A2916801), 1 mM sodium pyruvate (Gibco, 11360070), and 1.2 mM HEPES (Gibco, 15630080) for three passages following thawing from cryogenic storage. A total of 3 × 10⁶ cells were seeded in a 150 cm² culture flask containing 50 mL of complete DMEM. After seven days, the medium was replaced with an additional 50 mL of fresh DMEM and maintained for another seven days. The resulting L929-conditioned medium was then filtered using a 0.22 µm filter, aliquoted into 50 mL Falcon tubes, and stored at −20°C until use.

### Generation of bone marrow-derived macrophages

Bone marrow-derived macrophages (BMMs) were generated from 8- to 12-week-old male or female C57BL/6J mice by harvesting tibia and femur bones. Whole bone marrow (BM) cells were cultured in DMEM supplemented with 20% L929 cell-conditioned medium, 2 mM L-glutamine, 1 mM sodium pyruvate, 1.2 mM HEPES, 10% heat-inactivated FBS, and 1 U/mL penicillin/streptomycin (Gibco, 15070063). Cells were maintained in Petri dishes for seven days, with an additional 5 mL of supplemented medium added on Day 5. On Day 7, non-adherent cells were removed by washing with sterile PBS, and adherent macrophages were used for experiments. In a separate set of experiments, BMMs were generated by culturing whole bone marrow cells in the presence of recombinant murine M-CSF (30 ng/mL) (PeproTech, 31502).

### Antibodies and fluorescent conjugates

The following antibodies were purchased from BD Biosciences: anti-CD11b (clone M1/70) PE, 557397; anti-CD11c (clone HL3) PE, 557401, BV480, 565627 ; anti-CD40 (clone 3/23) PE, 553791; anti-CD44 (clone IM7) APC, 553991; anti-CD45 (clone 30-F11) APC and PercP, 559864, 557235 ; anti-I-A/I-E (clone 2G9) FITC, 553623; anti-Ly6G (clone 1A8) PerCPp-Cy5.5, 560602; anti-CD107a (clone 1D4B) BV421 and FITC, 564347, 553793 ; anti-CD107b (clone M3/84) FITC, 562061; anti-TNF (clone MP6-XT22) PE, 554419; anti-TREM-1(clone 174031) BV421, 747899 and Fixable Viability Staining 660, 564405. The following antibody was purchased from Thermo Fisher: anti-CD115 (clone AFS98) PercP 710, 46115282; The following antibodies were purchased from BioLegend: anti-CD11b (clone M1/70) BV421, 101251; anti-CD11c (clone N418) PerCP, 117325; anti-CD64 (clone X54-5/7.1) FITC, 139316; anti-CD68 (clone FA-11) PE, 137013; anti-CD69 (clone H1-2F3) PerCP Cy5.5, 104522; anti-CD74 (clone) Alexa Fluor 488, 151005; anti-CD88 (clone 20/70) PECy7, 135810; anti-CD107a (clone 1D4B) PECy7, 121619; anti-CD138 (clone 281-2) APC, 142506; anti-CD163 (clone S15049) PE,155307; anti-CD170 (clone S17007L) APC-Cy7,155531; anti-CD182 (clone TG11/CXCR2), 129103, anti-CD195 (HM-CCR5) APC and PE, 107012, 107006; anti-CD206 (clone C068C2) FITC, 141704; anti-CD207 (clone 4C7) PE-Cy7, 144209; anti-F4/80 (clone BM8) BV421, 123132; anti-Ly6C (clone HK1.4) Alexa fluor 488; 128021; and Zombie NIR, 423105).

### Flow cytometry and cell sorting and gating strategy

BMMs were detached from plates using accutase (Sigma-Aldrich, 6964), washed once with PBS, and resuspended in FACS buffer (PBS + 1% FBS). Samples were pre-incubated with Fc block (BD Bioscience, 552141) for 10 minutes on ice to minimize nonspecific binding. FACS sorting was performed using a Melody or ARIA II sorter with a 100-µm nozzle (Becton Dickinson). The gating strategy included exclusion of dead cells using Zombie NIR staining, selection of leukocytes as CD45⁺ cells, and subsequent gating based on forward scatter (FSC) versus side scatter (SSC) parameters. Within the final gates, the median fluorescence intensity (MFI) of F4/80 and CD11b was measured. Flow cytometry data were acquired using a FACSVerse (Becton Dickinson) or a Cytek® Northern Lights flow cytometer, and data analysis was performed with FlowJo software (version 10.8.1).

### RNA isolation and sequencing

Total RNA of sorted SSC^hi^FSC^hi^F4/80^hi^CD11b^hi^ and SSC^lo^FSC^lo^F4/80^lo^CD11b^lo^ macrophages, derived from three independent cell cultures, was extracted using TRIzol LS (Invitrogen; 10296010) treated with DNase I (Qiagen; 79254), and purified with the RNeasy Plus Micro kit (Qiagen; 74034). RNA quality was assessed using Pico Bioanalyser chips (Agilent, 5067-1513). Only RNAs displaying integrity numbers of 9 or higher were used for downstream procedures. RNAseq libraries were prepared using the Nugen Ovation Trio low input RNA Library Systems V2 (Nugen, 0507-08) according to the manufacturer’s instructions by the RU Genome Technology Center. Pooled libraries were sequenced as 50-nucleotide, paired-end reads on an Illumina HiSeq 2500 using v4 chemistry.

### RNAseq data quality assessment and visualization

Illumina sequencing adapters and reads with Phred quality scores <20 were removed with Trimmomatic. Trimmed reads were mapped to the *Mus musculus* genome Ensembl annotation release 91 using STAR v2.5.3a with default settings. The number of reads uniquely mapping to each gene feature in the corresponding annotation file was determined using featureCounts. The resulting count tables were passed to R for further analyses. RNAseq data are deposited at the Gene Expression Omnibus repository under accession #PRJNA1294819.

### Differential expression and gene ontology analyses

Consistency between replicates was checked using PCA and Euclidean distance–based hierarchical clustering on normalized counts. Weakly expressed genes, defined as having < 3 fragment per kilobase of transcript per million mapped reads in at least one group of replicates, were removed from differential expression analysis. Differential expression analyses were performed using the DESeq2 package. Genes were considered significant when the adjusted p-value was < 0.05. The set of significant genes for each contrast of interest was passed to enrichR and ClusterProfiler for functional gene enrichment analyses and heatmaps were performed using Pheatmap.

### Phagocytosis experiments

For flow cytometry-based quantification of mycobacterial phagocytosis by BMMs, cells were infected with SYTO24-stained Mtb or BCG at different multiplicity of infection (MOI). Unlabeled Mtb or BCG were used as controls for flow cytometry experiments. At different time points post-infection, cells were extensively washed with cold PBS and stained with antibody panels for flow cytometry analysis. In a separate set of experiments, BMMs were incubated with 50 µg/ml FITC-conjugated zymosan (Invitrogen, Z2841) for 4 hours or *Salmonella*-SYTO24 (MOI 10) for 30 min at 37°C, followed by extensive PBS washing before flow cytometry analysis. In a subset of experiments, cytochalasin D (Sigma-Aldrich, C8273 for Fig. 2 and Invitrogen, PHZ1063 for Supplementary Fig. 1B) was added 30 minutes prior to exposure to SYTO24-stained *M. bovis* BCG, FITC-conjugated zymosan or *Salmonella*-SYTO24. Cells were then incubated with bacteria for 4 hours at 37°C, followed by extensive PBS washing before flow cytometry analysis. To quantify mycobacterial uptake, MFI and frequency of positive macrophages were obtained by gating on specific cell subpopulations. Gating strategy was performed using controls of macrophage exposed to unlabeled particles. For microscopy-based quantification of mycobacterial uptake, BMMs were isolated from WT mice and allowed to adhere overnight onto poly-L-lysine-coated glass coverslips (Molecular Probes). Cells were then infected with SYTO24-stained Mtb at a MOI of 10 for 4 hours, followed by extensive wash and fixation with ProLong Gold antifade reagent containing DAPI (Invitrogen, P36935). The number of Mtb-infected cells was quantified across 50 fields (averaging 2 bacilli per cell) using a 100× objective on a BX41 microscope (Olympus Life Science). Data were expressed as the cell area versus the number of bacilli/cells.

### Cytokine production measurements

BMMs were seeded in 96-well flat-bottom plates at a density of 1×10⁶ cells/mL and infected with Mtb at a MOI of 3 for 4 hours. Following extensive washing to remove extracellular bacteria, cells were cultured for different time points. TNF, IL-6 and IL-10 were quantified in culture supernatants by commercially available ELISA. In separate experiments, BMMs were stimulated with Pam3Cys (S-[2,3-bis(palmitoyloxy)-(2-RS)-propyl]-*N*-palmitoyl-(R)-Cys-(S)-Ser-Lys4-OH, trihydrochloride) (100 ng/mL, EMC Microcollections), ultra-pure LPS (*E. coli* 0111:B4, Invivogen, tlrl-3pelps) (100 ng/mL), flagellin (100 ng/mL, Invivogen, tlrl-epstfla-5), or heat-killed BCG (MOI 10) for 6 hours at 37°C, with brefeldin A (BD GolgiPlug™, 555029) added during the final 4 hours. For intracellular TNF staining, single-cell suspensions were surface-stained, fixed, and permeabilized. Intracellular TNF was detected using anti-TNF monoclonal antibody (clone MP6-XT22). The percentage of TNF^+^ macrophages was determined by gating on specific cell subpopulations.

### Mtb killing assays

BMMs were seeded in 96-well flat-bottom plates at a density of 1×10⁶ cells/mL and infected with Mtb at a MOI of 3 for 4 hours. Following extensive washing to remove extracellular bacteria, infected macrophages were treated with recombinant murine IFN-γ (BD Pharmingen, 554587). At 24 hours and 48 hours post-infection, cells were washed and lysed with 1% saponin. Bacterial burden was assessed by plating lysates onto 7H10 Middlebrook agar for CFU enumeration. Nitrite production was measured using the Griess reaction (de Almeida et al, 2011), with absorbance recorded at 540 nm.

### In vivo infection model and flow cytometry experiments

C57BL/6J mice (8–12 weeks old) were housed under specific pathogen-free conditions at the animal facility of the Pharmacology Department, Universidade Federal de Santa Catarina. For infection, *M. bovis* BCG was suspended in PBS, and bacterial clumps were dissociated using a sterile needle attached to a syringe by repeatedly passing the suspension through the needle. Mice were infected via the intratracheal (i.t.) route with a total of 10⁸ CFU in PBS (50 µL) using a 0.25 mm needle gauge. At 18 h post-infection, whole lungs were collected and enzymatically digested with Liberase CI (Roche, 135476000) for 20 minutes at 37°C. Single-cell suspensions were prepared by passing the tissue through 100 µm cell strainers, and 2×10⁶ cells were used for flow cytometry staining. Gating strategy: Dead cells were excluded using Zombie NIR viability staining, and leukocytes were identified by gating on CD45^+^ cells. Within the CD45^+^ population, F4/80^+^CD11b^lo^ and F4/80^+^CD11b^hi^ macrophage subsets were defined. Alveolar macrophages (AMs) were identified as Siglec-F^hi^ cells within the F4/80^+^CD11b^lo^ gate. AM subsets were further characterized by CD107a and CD195 expression, delineating AM1 and AM2 populations.

### Statistical analyses

Statistical analysis was performed using GraphPad Prism software (version 10). The Shapiro-Wilk test was used to assess normality of the data. For comparisons between two groups, unpaired or paired Student’s t-tests were applied when data were normally distributed; when normality was not met (Shapiro-Wilk, p < 0.05), the Mann-Whitney U test (unpaired) or Wilcoxon matched-pairs signed rank test (paired) was used. For comparisons involving more than two groups or conditions, one-way ANOVA or two-way ANOVA was performed, followed by Tukey’s post hoc test when data passed normality. Statistical significance was defined as p < 0.05 (two-tailed). To evaluate the magnitude and biological relevance of the findings, Cohen’s d was calculated as a standardized effect size, and 95% confidence intervals were computed for mean/median differences and fold changes where applicable.

## Acknowledgments

We express our gratitude to Prof. Juliano Ferreira (UFSC) for providing animal housing facilities, Prof. Dario Zamboni (FMRP-USP) for supplying BMs from CD195-deficient mice, and Prof. Daniel Mucida (Rockefeller University) for granting access to RNA sequencing facilities. We sincerely thank Dr. Alan Sher (National Institutes of Health) for his insightful and constructive comments. This work was funded by Howard Hughes Medical 802 Institute – Early Career Scientist (AB; 55007412), and CNPQ/PQ Scholars (ALBB, DSM and ELR).

## Author Contributions

CE, HMS and ALBB designed research; CE, GL, HMS, ELM, DAGBM, BKB, LZM, MCCC, BSR, MS, GS and JB performed research; YKTM, DSM and ELR contributed new reagents/analytic tools; CE, GL, HMS, MS, ELR and ALBB analyzed data; and CE and ALBB wrote the paper.

---

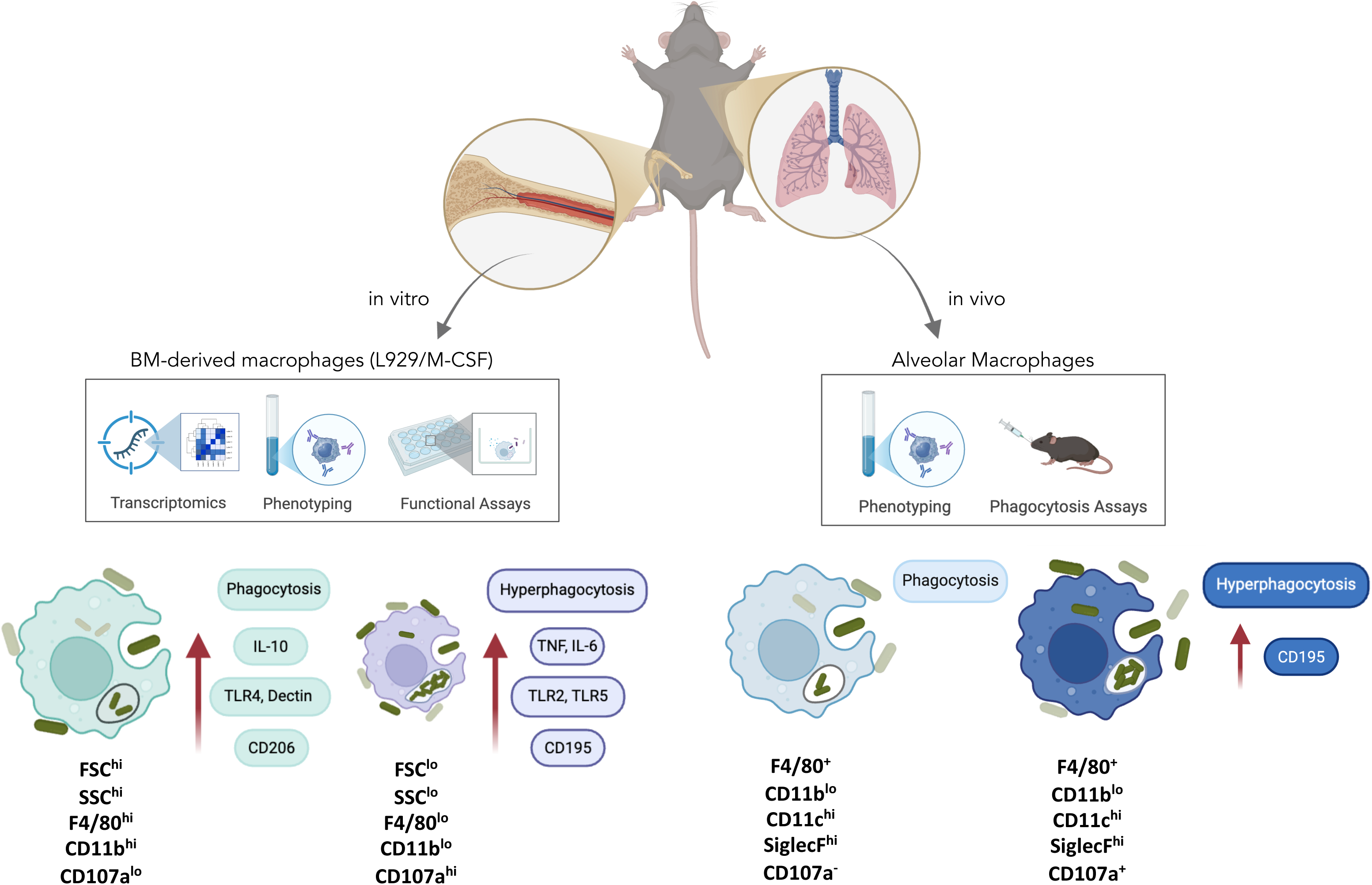

**Supplementary Figure 1.**
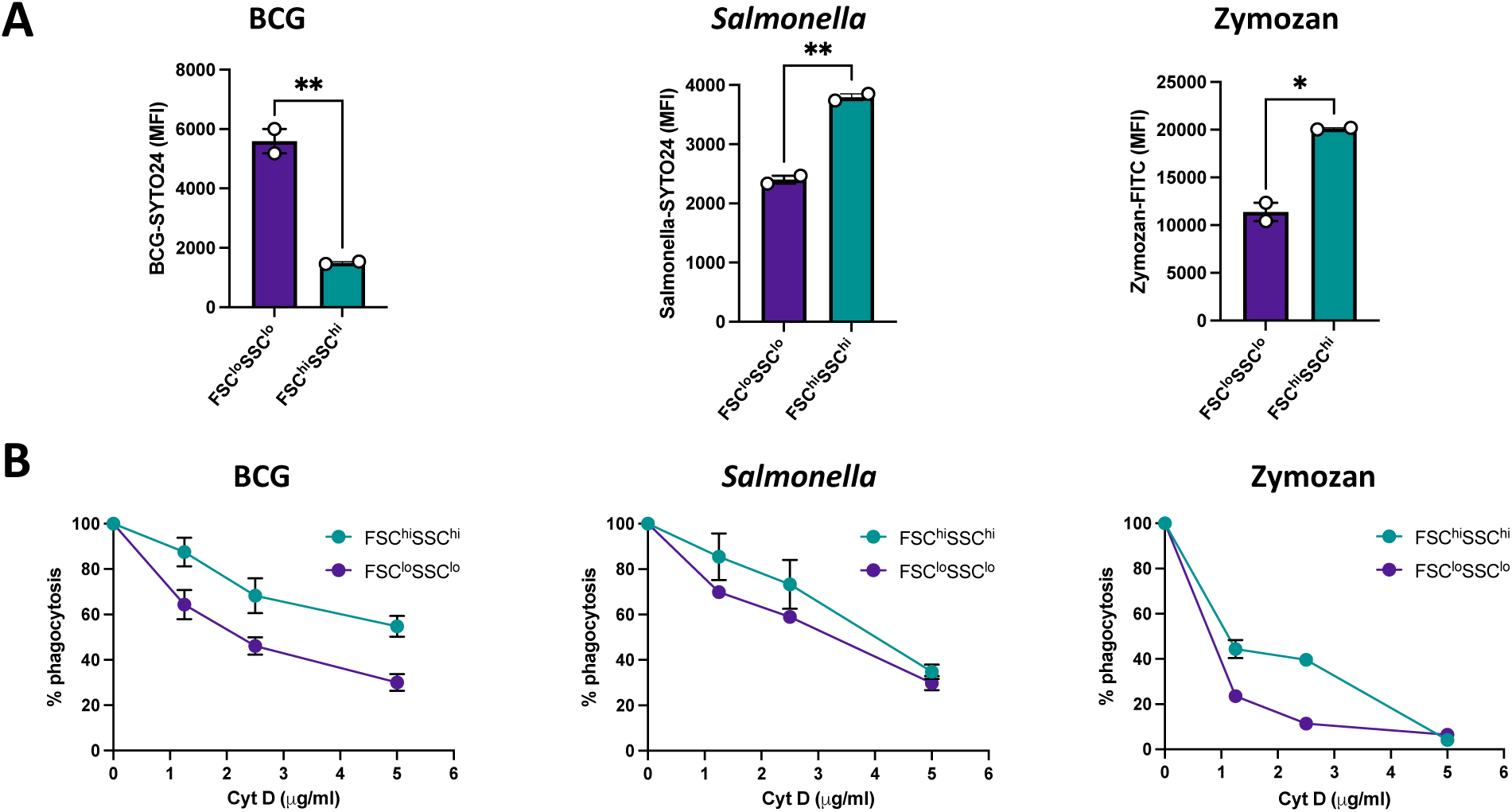
Cytochalasin (Cyt D) differentially inhibits phagocytosis of BCG, *Salmonella* and zymosan by FSC^hi^SSC^hi^ and FSC^lo^SSC^lo^ macrophages. Bone marrow–derived macrophages (BMMs) were pretreated with cytochalasin D (1, 2.5, or 5 μg/mL) for 30 min prior to exposure to BCG-SYTO24 (MOI 10, 4 h), *Salmonella*-SYTO24 (MOI 10, 30 min), or zymosan-FITC (50 μg/mL, 4 h), followed by flow cytometry analysis. **(A)** Mean fluorescence intensity (MFI) of BCG, *Salmonella*, or zymosan uptake in FSC^hi^SSC^hi^ and FSC^lo^SSC^lo^ macrophages. **(B)** Dose–response effect of cytochalasin D (1, 2.5, or 5 μg/mL) on phagocytosis by FSC^hi^SSC^hi^ and FSC^lo^SSC^lo^ macrophages. Data represent mean ± SEM from two independent mice.

